# Cut homeodomain transcription factor is a novel regulator of cortex glia morphogenesis and maintenance of neural niche

**DOI:** 10.1101/2022.10.31.514621

**Authors:** Vaishali Yadav, Ramkrishna Mishra, Papri Das, Richa Arya

## Abstract

Cortex glia in *Drosophila* central nervous system forms a niche around neural cells for necessary signals to establish cross-talk with their surroundings. These cells grow and expand their thin processes around neural cell bodies. Although essential for the development and function of the nervous system, how these cells make extensive and intricate connected networks remain largely unknown. Here we show that Cut, a homeodomain transcription factor, directly regulates the fate of the cortex glia, impacting NSC homeostasis. Focusing on the thoracic ventral nerve cord (tVNC), we found that Cut is required for normal growth and development of cortex glia and timely increase in DNA content to undergo endomitosis. Knockdown of Cut in cortex glia significantly reduces the growth of cellular processes, the network around NSCs, and their progeny’s cell bodies. Conversely, overexpression of Cut induces overall growth of the main processes at the expense of side ones. Whereas the Cut knockdown slowdown the timely increase of DNA, Cut overexpression results in a significant increase in nuclear size and volume and a threefold increase in DNA content of cortex glia. Further, we note that constitutively high Cut also interfered with nuclei separation during endomitosis. Since cortex glia form syncytial networks around neural cells, the finding identifies Cut as a novel regulator of glial growth and endomitosis to support a functional nervous system.

**Article Summary:** Cut homeodomain transcription factor is crucial for cortex glia growth and the formation of complex cellular processes around neural cells. This regulation ensures a timely increase in DNA content, allowing the cells to enter endomitosis. Constitutively high Cut levels increase the DNA content of these cells to several folds. The finding emphasizes the need to investigate if activated CUX1, the human homolog of Cut, in glioma enhances chromosomal instability and, in conjunction with other mutations, enhances their tumorigenic potential.

## Introduction

Glia cells play a crucial role in maintaining and protecting the nervous system by promoting neural growth, facilitating communication, and providing nourishment and support for the overall homeostasis of neural stem cells (NSC) and their progeny (Xiong and Montell 1995; Booth et al. 2000; Barres 2008; Trapp and Nave 2008; Stork et al. 2012). In addition, these cells form the blood-brain barrier and compartmentalize the central nervous system (CNS) into specialized domains in several invertebrates and vertebrates such as elasmobranch fishes(Oland and Tolbert 2003; Abbott 2005; Awasaki et al. 2008). It is particularly interesting to understand how glial cells form a dynamic environment around NSCs and their offspring during normal development and disease. Mammalian and *Drosophila* glial cells are comparable and classified into functional groups based on morphology and molecular functions (Stork et al. 2012). In Drosophila, the three major classes of glia types based on morphology are surface, cortex, and neuropile-associated glia (Ito, Urban, Technau, et al. 1995; Stork et al. 2012).

Cortex glia cells in *Drosophila* nervous system are born during the mid-embryonic stage. Their cell bodies are irregular and grow in multiple directions making massive networks around individual neural cell bodies for trophic and metabolic support (Ito, Urban, Technau, et al. 1995; Dumstrei et al. 2003; Pereanu et al. 2005). In the ventral nerve cord (VNC) of the *Drosophila* larval nervous system, cortex glia are primarily located on the ventral and lateral sides. These cells closely interact with neural cells and also play a role in regulating their fate (Ito, Urban, and Technau 1995; Renee D Read 2018; Dong et al. 2021). Similar to cortex glia, mammalian astrocytes also extend their processes toward the outer surface of synaptic neuropils (reviewed in Zhou et al. 2019). In order for glial processes to grow, Insulin and PI3K signaling must be activated through the intake of nutrients (Spéder and Brand 2018; Yuan, Conor W. Sipe, et al. 2020). In addition, Hippo, EGFR, and FGF signaling pathways also regulate cortex glia growth and remodeling (Read et al. 2009; Witte et al. 2009; Reddy B. V. V. G. and Irvine Kenneth 2011; Avet-Rochex et al. 2012; Spé Der and Brand 2018; Dong et al. 2020). Investigations continue how fine processes grow from the cortex glia cell body and find their correct path. We discovered that Cut, a homeodomain transcription factor, is critical and play multiple roles in the developmental growth of cortex glia to form a niche around neural cell bodies.

The Cut protein family is abundantly expressed in the developing central nervous system of both *Drosophila* and mammals, playing a crucial role in regulating neural identity and proper patterning. (Arya et al. 2019; Weiss et al. 2019). From an evolutionary and functional perspective, Cut is one of the most conserved proteins among all metazoans. It is known as CUX1 in humans and CUX1/2 in mice (Neufeld et al. 1992; Valarche et al. 1993; Yoon and Chikaraishi 1994; Quaggin et al. 1996; Tavares et al. 2000). Cut protein is well known for its diverse functions in determining cell specificity and identity in *Drosophila* and mammals and is expressed in multiple tissues (Liu et al. 1991; Andres et al. 1992; Liu and Jack 1992; Ludlow et al. 1996; Nepveu 2001; Zhai et al. 2012; Pitsouli and Perrimon 2013). It influences cell proliferation, migration, and response to DNA damage in various tissues (Coqueret et al. 1998; Michl et al. 2005; Cadieux et al. 2009; Vadnais et al. 2012; Pitsouli and Perrimon 2013) The protein Cut has multiple DNA binding domains, resulting in various transcription factor isoforms (Bodmer et al. 1987; Blochlinger et al. 1988; Liu et al. 1991). Its isoforms have a single homeodomain and one or multiple Cut DNA binding repeat domains. Depending on its interacting partners, Cut acts as a transcriptional activator or repressor of several genes during development (Yoon and Chikaraishi 1994; van Wijnen et al. 1996).

Consistent with its role as a gene regulator, Cut is also identified as a selector gene in several mammalian disorders, including cancer (Zhai et al. 2012). Loss of function Cut/CUX1 mutations have been identified in several cancer types. At the same time, increases in CUX1 gene copy number and overexpression are also common in many cancers (Sansregret and Nepveu 2008; Ramdzan and Nepveu 2014). In the mouse, CUX1 overexpression leads to multiorgan hyperplasia and tumor development after a long latency period (Ledford et al. 2002; vanden Heuvel et al. 2005). Since Cut perform a diverse function in several cellular processes and gene regulation, the detailed molecular understanding of its functional diversity is an area of active research.

This research reports that the Cut homeodomain transcription factor is crucial in regulating the growth and formation of the cortex glia network around NSCs and their progeny in the *Drosophila* central nervous system. Cut guides the cortex glia development in several ways. Cut levels affect the complex network of glial processes. Loss of Cut does not allow cortex glia to undergo timely endomitosis to increase their nuclei number. On the other hand, continued high Cut expression in cortex glia increases their DNA ploidy to multifold, yet it also interferes with the normal endomitosis by not allowing the nuclei to split indicating a gain of function for Cut. It is the first report highlighting a novel function of Cut in regulating the growth and branching of cortex glia and control of endomitosis.

## Results

### 1. Cut expresses in glial subtypes and maintains niche around neural stem cells

Transcription factor Cut is known to express in *Drosophila* CNS and to regulate the developmental death of NSCs in the abdominal region of VNC (Arya et al. 2019). In addition to NSCs, Cut is expressed in several other cells in the larval CNS. Therefore, the larval CNS was stained with a Cut antibody to profile its expression in the glial cells. In this study, we focussed on the thoracic region of the ventral nerve cord (tVNC) to study the role of Cut in glial biogenesis (Fig. 1A). Since Cut and pan-glia marker Repo antibodies are both raised in mice, and there is no commercial antibody for cortex glia, we used the *UAS-GAL4* system to mark the glia with GFP using *Repo-GAL4* (pan glia) and *Cyp4g15-GAL4* (cortex glia) drivers (Fig. 1B, C, Fig. S1 D-I) (Spéder and Brand 2018; Gonzalez-Gutierrez et al. 2019 Apr; Rujano et al. 2022). We counted the GFP+ and Cut+ nuclei in the tVNC to estimate the glia that express Cut. In *Repo>GFP* tVNC, where GFP marks all the glial cells, around 55% of total GFP-positive nuclei were Cut positive (Fig.1B, graph G, Fig. S1. D-F). On the other hand, when the numbers of GFP+ nuclei marking the cortex glia *(Cyp4g15>GFP*) that also express Cut were counted, we found that all the tVNC cortex glia express Cut (Fig.1C, graph G, Fig. S1.G-I).

**Figure 1:**
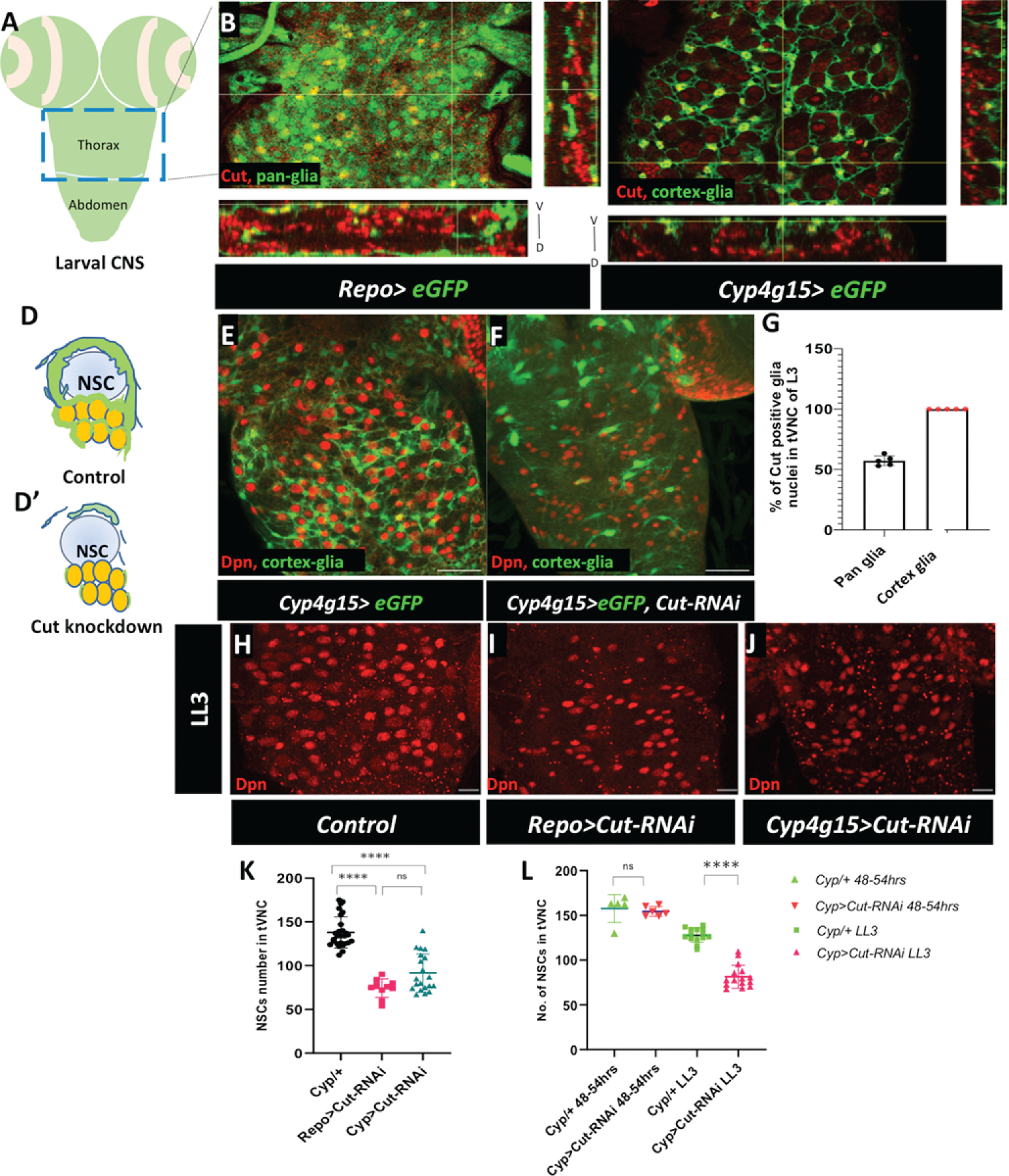
Cut expresses in glial subtypes and maintains niche around neural stem cells. **(A)** Model of larval CNS indicating the thoracic region and abdominal region of VNC. The blue box marks the late thoracic region of VNC (tVNC), shown in **(B-J)**. **(B-**C) Single section view and orthogonal sections in xyz, and yxz planes shows Cut expression (red) in pan glia (*Repo>eGFP*, **B)** and cortex glia (*Cyp4g15>eGFP,* **C). (D)** model showing cortex glia (green) enwrapping NSC (sky blue) and its lineages (yellow) in control and **(D’)** disruption of the cortex glia network in Cut-knockdown. **(E)** well-developed glial extensions around NSCs (marked with Dpn in red) in control (*Cyp4g15>eGFP/+),* **(F)** Loss of Cut (*Cyp4g15>eGFP, Cut-RNAi*) severely disrupts cortex glia and affects the distribution of NSCs. **(G)** Quantification of the percentage of Cut positive glia nuclei in control tVNC. (H-J) shows NSC number and distribution (Dpn, red) in tVNC **(H)** control (*Repo-GAL4/+),* **(I)** Pan glial Cut knockdown (*Repo>Cut-RNAi)*, and (J) cortex glia Cut knockdown (*Cyp4g15>Cut-RNAi).* **(K)** Quantification of NSCs loss upon Cut-knockdown with *Repo-GAL4* and *Cyp4g15-GAL4* compared to control. **(L)** Quantification in ALH 48-54 hr and LL3 (late larval stage 3) to estimate the non-autonomous loss of NSCs upon Cut-knockdown in cortex glia, Statistical evaluation of significance based on unpaired t-test is marked with stars, **** p<0.0001. Scale bar: 50µm (E, F), 20µm **(H-J).**

The cortex glia networks around neural cells, establish the communication between the neural cells to their environment and are needed for the survival of newborn neurons (Freeman and Doe 2001; Dumstrei et al. 2003; Spéder and Brand 2018; Rujano et al. 2022). Selective knockdown of Cut expression in cortex glia with *Cyp4g15-GAL4* driver resulted in substantial disruption of their cellular processes (compare Fig.1D’ with D and F with control E). We validated this finding with two separate *Cut-RNAi* lines (Fig.S1A, B). Since the cortex glia makes a supporting system for the neural cells, the loss of glia and their network around neural bodies also affects NSCs. We used Deadpan (Dpn) as an NSC marker and noted that they were unusually clumped, and were eliminated during development upon Cut ablation in cortex glia (Fig.1H-J, graph K). To ensure that the fewer NSCs observed at the late third instar larvae are not due to an early defect in their birth, we counted NSCs in the early instar stages. The number of NSC in tVNC of control and Cut knockdown did not show any significant difference in early third instar larvae (Fig.1L), indicating that their number was normal initially and declined only later due to loss of the niche (Fig.1L). Thus, Cut defective cortex glia non-autonomously affects the NSC fate. We used four other cortex glia-specific drivers for knockdown of Cut - *R54H02, R46H12, and Np577* and found similar, significant non-autonomous loss of NSCs in tVNC (Fig.S1C). We selected *Cyp4g15-GAL4* for further studies. We conclude that Cut is expressed in all cortex glia and is essential for maintaining the glial niche around the NSCs and their progeny.

### 2. Cut is required for the growth of the cortex glia processes and their branching

Following the above result that Cut defective cortex glia cannot form a niche around the neural cells, we further explored the role of Cut in these cortex glia for their growth and chamber formation around neural cell bodies. Cortex glia continues to expand their cytoplasmic extensions from the late embryonic stage (Pereanu et al. 2005; Jaeda C. Coutinho-Budd et al. 2017; Rujano et al. 2022). Therefore, as a first step, we checked the expression profile of the cortex glia GAL4 driver, *Cyp4g15-GAL4*, and found that it expresses from the late embryonic stage (Fig. S2 I, I’) and continues its expression through different larval stages. The GFP driven with the GAL4 line faithfully marks the glial network from the early to late developmental stages of larvae (Fig. S2 I, I’ with Repo, Fig. 2A, 3A-H). As shown in Fig. 2A, C (yellow arrowhead in 2C), multiple thicker and thin cytoplasmic extensions emerge from a cortex glia cell body and spread their branches in multiple directions making a fine meshwork around individual neural cell bodies in the tVNC, known as trophospongium (Fig 2C, red arrow). Interestingly, the cellular processes from different cortex glia are well connected and self-tilled to one another (Jaeda C. Coutinho-Budd et al. 2017; Rujano et al. 2022). Cut knockdown in the cortex glia leads to a massive loss of these lamelliform cortex processes in the whole CNS (Fig. 2B-B’’, Fig.S2E-F’’for optic lobes and S2G-H’, graph J for the abdominal region). Unlike the normal cortex glia, which extends the processes in several directions (Fig. 2C yellow arrowheads), the Cut knockdown in these cortex glia leads to only one to two abnormal-looking main cytoplasmic extensions (compare Fig. 2C, D). To further compare the density of cortex glia extensions in the two genotypes, we used the ImageJ plot analysis tool to evaluate the number of peaks, indicating the number of cortex glia extensions. Cut defective cortex glia show far less peaks than the control. Additionally, the intensities of peaks were also variable, indicating a reduction in cytoplasmic extensions (Fig S2C, D). The plot profile along a line (Fig S2A, B’) in the tVNC also showed a loss of the glial network upon Cut knockdown (Fig. 2E, F). We also evaluated the volume of cortex glia in these genotypes through thresholding and network density through ImageJ, where the volume of Cut defective cortex glia is significantly reduced.

**Figure 2:**
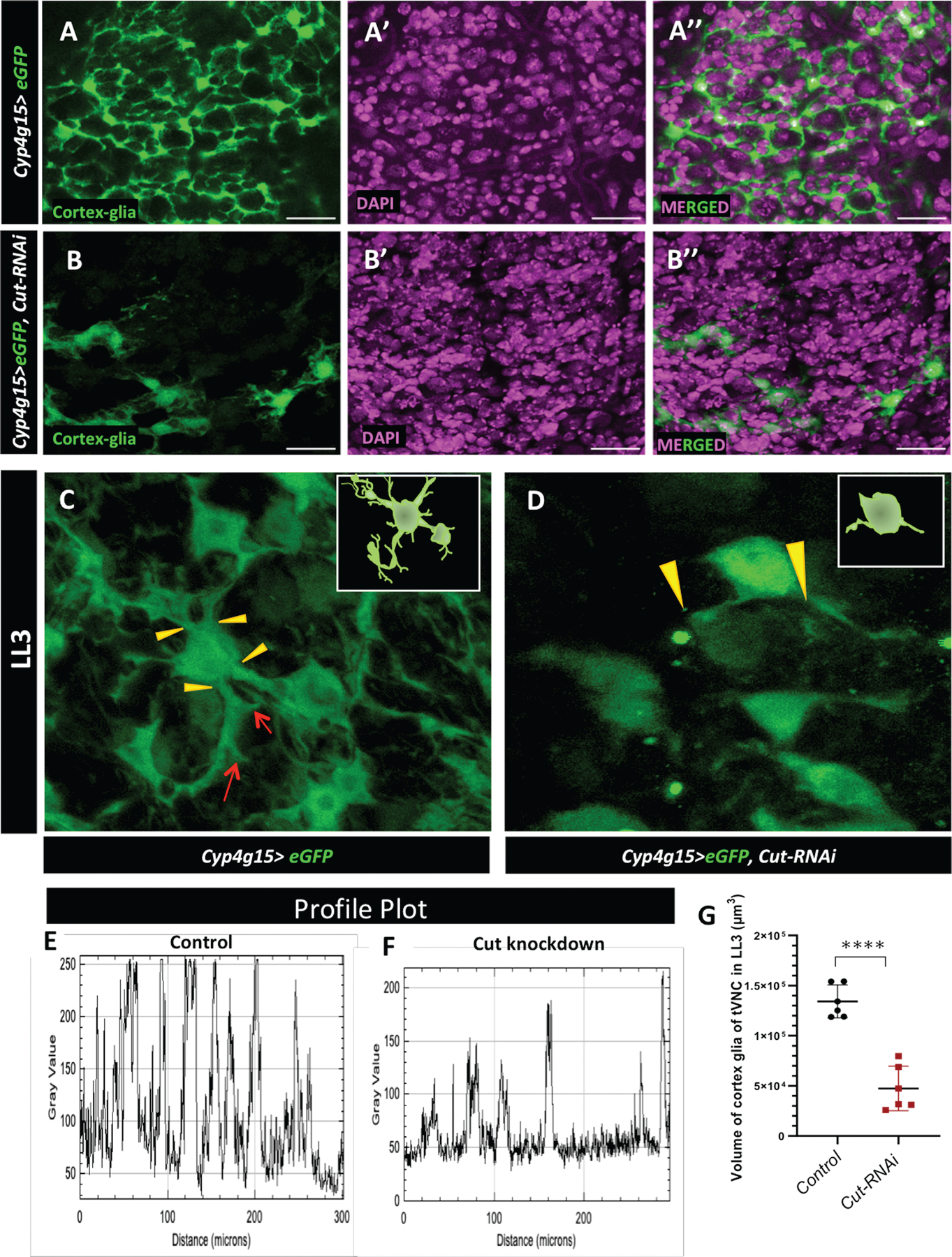
Cut is required for cortex glia growth and branching. **(A-A’’)** Control *(Cyp4g15>eGFP/+)* shows cortex glia niche enwrap the neural cells. **(B-B’’)** Cut defective cortex glia cannot form a niche around the neural cell bodies (green cortex glia, magenta DAPI). **(C)** In Control *(Cyp4g15>eGFP/+)* individual cortex glia cell body extends thick (yellow arrowheads) and lean processes (red arrows) in multiple directions; inset is the model. **(D)** Cut defective cortex glia (*Cyp4g15>eGFP, Cut-RNAi*) with minimum extension growth. **(E, F)** plot profile along a straight line in tVNC showing that peaks and their intensities which are more uniform and closer to each other in control **(E)** compared to Cut knockdown **(F),** emphasizing the loss of glial network. **(G)** Evaluation of the Volume (µm^3^) of cortex glia in LL3 tVNC of control *(Cyp4g15>eGFP/+*), and Cut Knockdown *(Cyp4g15>eGFP, Cut-RNAi*) showing significantly reduced cortex glia volume upon Cut-Knockdown, Statistical evaluation of significance. ****p<0.0001, based on unpaired t-test. Scale bar: 20 µm

The finding that Cut knockdown prevents cortex glia nuclei from growing extensions and filling gaps, is significant since these cells normally grow efficiently when a few of their neighbors are ablated (Jaeda C. Coutinho-Budd et al. 2017). This indicates that Cut is required for the normal growth and development of lamelliform extensions from the cell body and that upon Cut knockdown, several of these defective glia could not undergo proper morphogenesis.

### 3. Cut defective cortex glia are unable to increase their nuclei number during development

To better understand the defect of cortex glia in tVNC during development and its time of onset, we examined CNS at different time points during larval stages. In the early first larval instar (L1, ALH 0-3hr), there is typically 4-5 cortex glia in each thoracic hemisegment arranged in a ventrolateral pattern (Ito, Urban, Technau, et al. 1995, Ito et al 1995). One cortex gila can be seen near the midline, two are placed ventrally, and one or two are placed laterally (Fig. 3A, A’). At this stage, the extensions coming out of the cortex glia are very fine. In L2 (ALH 24-28hr), the extensions are more distinctly visible and infiltrate throughout tVNC. However, the number of cortex glial nuclei remains largely unchanged compared to L1 (compare Fig. 3C, C’ with A, A’) (Ito, Urban, and Technau 1995). Cut knockdown in cortex glia impacts the networking of cellular processes and nuclei number as early as ALH 24-28hr (fig. 3D, D’). Even though the number of cortex glia nuclei does not show severe change compared to L1 of Cut knockdown, the connection of cellular processes is severely affected, and fine networks are mostly missing at this point (Fig. 3D, graph I arrow).

**Figure 3:**
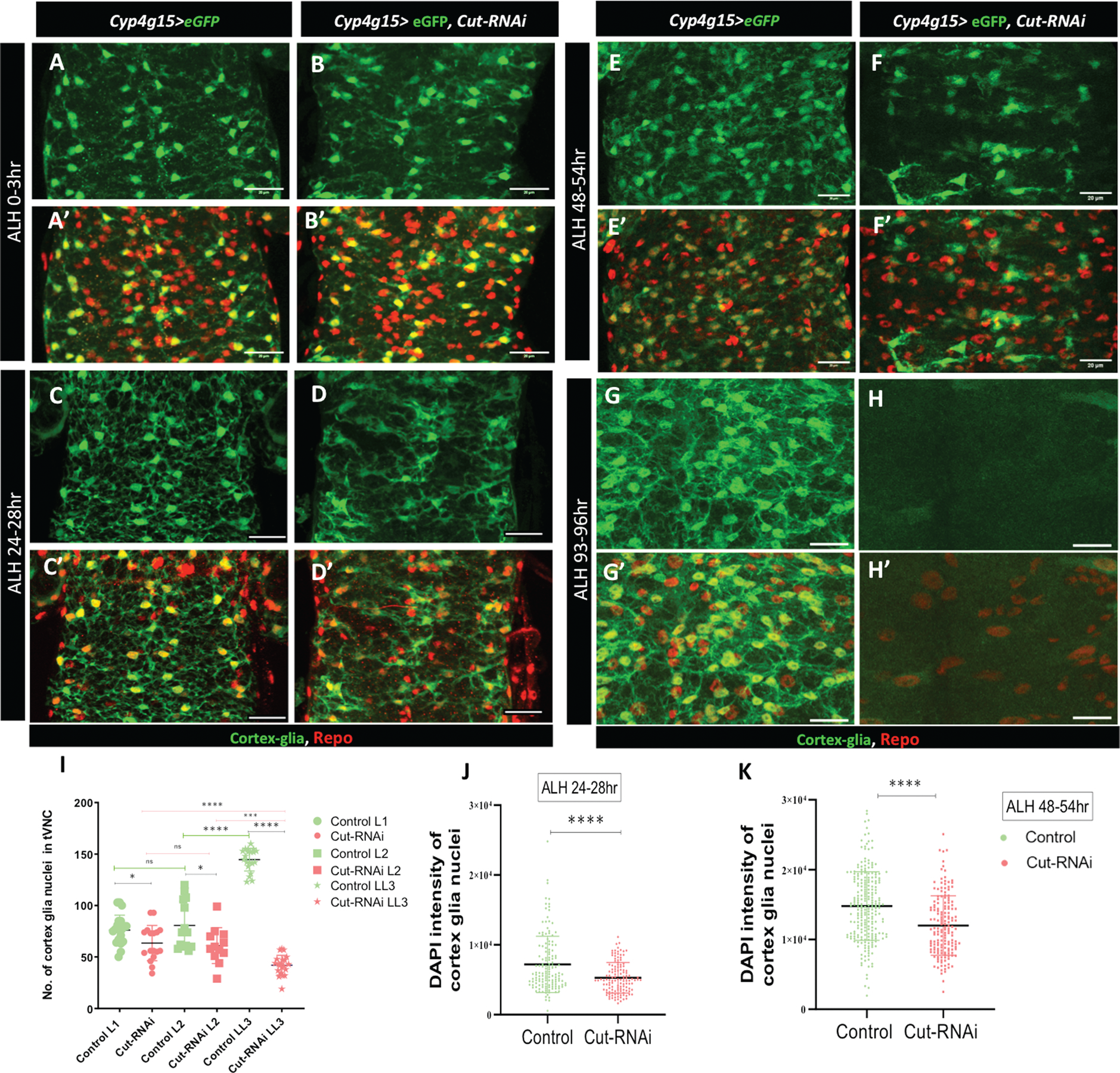
Cut defective cortex glia are unable to increase their nuclei number during development. **(A-K)** Throughout the figure the genotypes of control (*Cyp4g15>eGFP/+)* and Cut knockdown (*Cyp4g15>eGFP, Cut-RNAi*), **(A-B’)** projection of ventral sections of only tVNC in early first larval instar (L1, ALH 0-3hr), in **(A, A’)** control and **(B, B’)** Cut knockdown 3-4 cortex glia are visible in each thoracic hemisegment, one near the midline and two ventral, lateral ones are not clearly visible in all hemisegments. **(C-D’)** L2 (ALH 24-28hr) in **(C, C’)** control, the cortex glial nuclei remain largely unchanged compared to L1 however the processes are more distinctly visible and infiltrate throughout tVNC, Upon Cut knockdown **(D, D’)** the cortex glia nuclei appear to be a little less and the cell processes are disrupted compared to control. **(E-F’)** L3 (ALH 48-54hr) in **(E, E’)** control the number of cortex glial nuclei increases significantly and the cell processes make a complex network, in **(F-F’)** Cut defective cortex glia shows defect in the increase in nuclei number, they still resemble the L1/L2 pattern of nuclei distribution (compare **F** with **A, C),** the glia process remains stunted. **(G-H’)** late L3 (ALH 93-96hr) in **(G, G’)** control the nuclei number further increases and the glia network is more like a honeycomb structure, upon Cut knockdown **(H, H’)** significant loss of cortex glia is noticed. **(I)** quantification of cortex glia nuclei in different larval timepoints (L1-L3), in L1 the number of nuclei in Cut defective cortex glia are slightly less compared to control (p value 0.0207). The nuclei number increases significantly in control from L2 to L3 (****p value<0.0001), upon Cut-knockdown the increase in nuclei number is largely hampered (*p value 0.0257) and in L3 several of them are lost (****p value <0.0001). **(J-K)** Quantification of the integrated density of DAPI of individual cortex glia nucleus at **(J)** ALH 24-28hr and **(K)** ALH48-54hr was evaluated (>5CNS/genotype). The CNS were stained with Repo (glia), GFP (cortex glia), DAPI (nucleus). **(J)** DAPI content of cortex glia is more, compared with Cut knockdown (****p value <0.0001) at ALH 24-28hr. **(K)** at ALH48-54 the DNA content is furthermore in cortex glia compared Cut defective cortex glia (****p value <0.0001). Statistical evaluation of significance based on unpaired t-test. Scale bar: 20 µm

The number of cortex glia nuclei typically increases from L2 onwards due to endomitosis (graph I) (Rujano et al. 2022). Furthermore, fusion of cortex glia processes results in the formation of a complex pattern of nuclei and cellular network (Jaeda C Coutinho-Budd et al. 2017; Rujano et al. 2022) (Fig. 3E-E’, graph I). Strikingly, the Cut knockdown affects the increase in nuclei number in L3 (48-96 hr) (compare Fig. 3E, E’ with F, F’, and Fig. 3G,G’ with H,H’). It is interesting to note that at this stage the nuclei pattern in Cut knockdown tVNC resembles more to the L1/L2 arrangement rather than L3 controls of the same age (compare Fig. 3F, F’, with Fig. 3C,C’ also 3 F,F’ and 3E,E’) the cell bodies appear flatten and unconnected.

Normally cortex glia shows a significant increase in nuclear volume from L2 onwards which is needed for endocycle and endomitosis (Rujano et al. 2022), we further investigate if Cut plays a role in DNA increase. We measured the DNA content through the integrated density of DAPI of each cortex glia nuclei labeled with GFP and Repo (DAPI+, GFP+, Repo+) using ImageJ. The nuclear content of cortex glia shows a sharp increase from ALH 24 to ALH 48hr as shown before (graph 3J, K). We observed that upon Cut knockdown, the integrated density of DAPI does not show the kind of increase as seen in the control group (graph 3J, K). This indicates that low Cut levels hinder the timely increase of DNA in cortex glia.

We further checked how Cut knockdown in the cortex glia affects an organism’s survival. We observed significant pupal lethality during development. However, we did notice a developmental delay progressing through different stages (Fig. S3A-B). In adult organisms, a decrease in Cut resulted in a significant reduction in average life span compared to control animals (Fig. S3B). Since Cut knockdown does not show severe consequences on the overall life span of the organism it may be due to a late and progressive effect on the loss of cortex glia morphogenesis. A very severe effect on glial growth upon Cut depletion occurs later in larval life (Fig. 3D-H), which may allow the organism to survive into adulthood after passing the critical stage of NSC activation during the larval life. We hypothesized that Cut is required for the timely growth of cortex glia, which allows them to undergo endomitosis and increase their nuclei number.

### 4. Overexpression of Cut enhances the growth of cortex glia main branches at the cost of side extensions

During *Drosophila* development, Cut influences the complexity of the neuronal dendritic arbor in the peripheral nervous system, determines the identity of wrapping glia, and influences the formation of cellular processes (Grueber et al. 2003; Bauke et al. 2015). Furthermore, the Cut protein level also regulates the branching pattern of dendrites in various neuronal classes in the peripheral nervous system differentially (Grueber et al. 2003). Therefore, we asked if constitutively increased Cut levels in cortex glia would affect their growth and branching patterns.

The cortex glial cells make a complex network of processes around the neural cells. These processes come in various lengths and thicknesses. Some of them originate directly from the cell body as the main branch, while others stem from the main branch as side processes with varying thicknesses. We found that the Cut in the cortex glia remarkably affects the cellular processes in several ways. While Cut knockdown resulted in stunted growth of cortex glia processes as noted above (Figs. 2, 3), its over-expression led to longer main extensions but with little side branches (see Fig. 4A, D, red arrows and yellow dotted lines marking main extensions). Note in control where one of the marked main extensions coming out of the cell body is shorter (Fig.4A, red arrow, yellow line); in Cut overexpressing glia it grows thicker and longer (Fig.4D, red arrow, yellow line). To validate the visual interpretation, we measured the total volume of cortex glia and found an apparent increase in overall glial volume upon Cut overexpression (Fig. 4H). Since Cut overexpressing glia has more extended processes, the cortex glia chambers are also larger and globular (compare Fig.4 B, C with E, F, graph G). However, many fewer thinner processes that enwrap the individual neural cell bodies are typically seen in the control tVNC (compare Fig. 4E with B). Further, to check the extent of loss of side fine processes upon Cut overexpression, we compare the density of main vs. side extensions through the surface and plot profile along a line. The loss of side extensions upon Cut overexpression is very evident as the profile of Cut overexpressing glia has broader peaks, away from each other, compared to the control, where the peaks are more uniform, sharp, and close to each other (Fig. 4I-L). Thus, these observations all together propose that high Cut levels are sufficient to influence the overall growth of cortex glial processes. In addition, the Cut level affects the glial network’s complexity by increasing the growth of main processes and retard the growth of side extensions.

**Figure 4:**
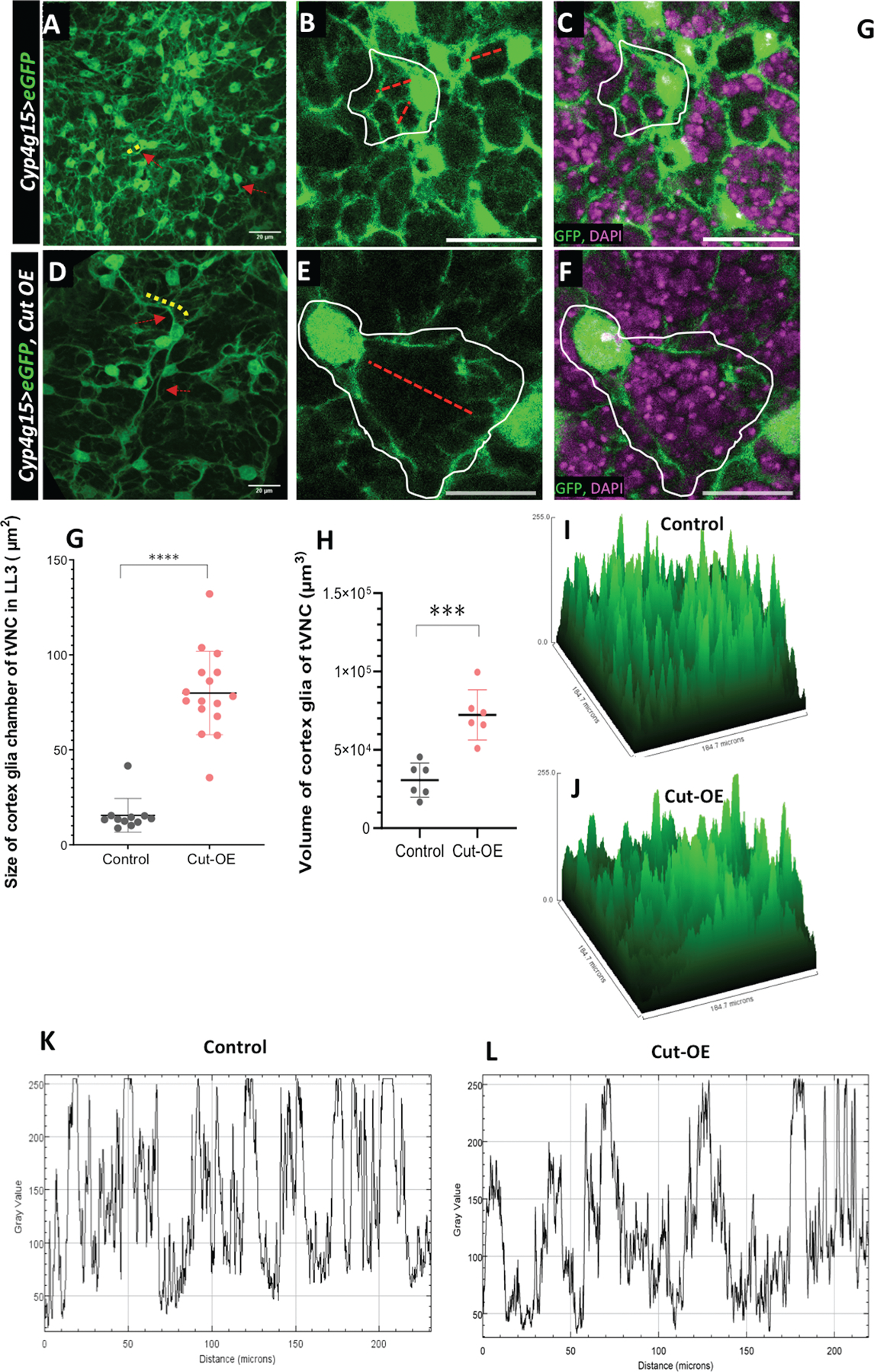
Overexpression of Cut in cortex glia enhances the growth of cell body and main branches at the cost of terminal extensions. **(A-C)** are control (*Cyp4g15>eGFP/+);* **(A)** cortex glia cell bodies (red arrows) and dense main extension (yellow lines) can be seen, (**B**) the cortex glia chambers (white encircle) are small and compact (red line to show the area covered), **(C)** same image in **(B)** with DAPI showing single DAPI neural nuclei in the encapsulation**. (D-F)** are Cut overexpressing cortex glia *(Cyp4g15>eGFP, Cut-OE),* Compare **D-F** with control **A-C** where cell bodies are less dense, big, and their cytoplasmic extensions are longer and thicker, the main cortex glia branches are significantly longer **(D**), making bigger glial chambers **(E),** and encapsulating several neural bodies marked with DAPI **(F)**. **(G-H)** quantifications in control *(Cyp4g15>eGFP/+)* and Cut overexpression *(Cyp4g15> eGFP, Cut-OE).* **(G)** size of the cortex glia chamber, total chambers counted 5/tVNC (n=3) µm^2^. **(H)** total cortex glia volume in tVNC (n>5). **(I,J)** represents the surface plot of cortex glia in tVNC to compare the intensity and extensions of Cut overexpressing cortex glia cells with control. In Control **(I)** there are more peaks representing cell and their extensions. In contrast, upon Cut-overexpression **(J)** the number of peaks is less and of variable intensities showing the loss of cortex glia side extensions. **(K,L)** is the plot profile along a line in the tVNC showing the peaks and their intensities in Control **(K)** and Cut overexpression **(L),** again emphasizing the loss of fine glial network. Statistical evaluation of significance ****p<0.0001, ***p<0.001, and **p<0.01 based on unpaired t-test. Scale bar: 20µm.

### 5. Constitutive activation of Cut increases DNA content in cortex glia

The number of cortex glia nuclei in the tVNC progressively increases from the L1 to LL3 stages due to endomitosis (Fig. 5A-C) (Unhavaithaya and Orr-Weaver 2012; Jaeda C Coutinho-Budd et al. 2017; Rujano et al. 2022). Cut overexpressing cortex glia in tVNC increases their nuclei number from the L1-late L2 stage, (compare Fig. 5D, E with 5A, B graph H). On the contrary, while this number increases further in control tVNC in the late L3 stage (Fig.5B-C, graph G), the rise is not seen in the Cut overexpressing cortex glia nuclei number (Fig.5E-F, graph G). Instead of an increase in the nuclei number, the cortex glia constitutively expressing Cut increase their nuclei size (compare Fig. 5graph H). We observe a threefold increase in nuclei size upon Cut overexpression (Fig. 5I). Furthermore, we observed that the distribution pattern of cortex glia nuclei in LL3 tVNC of Cut overexpressing CNS is more similar to the control L1/L2 tVNC instead of LL3 (compare Fig. 5F and 5A-B), although the pattern is deformed due to their unexpected loss (Fig. 5F).

**Figure 5.**
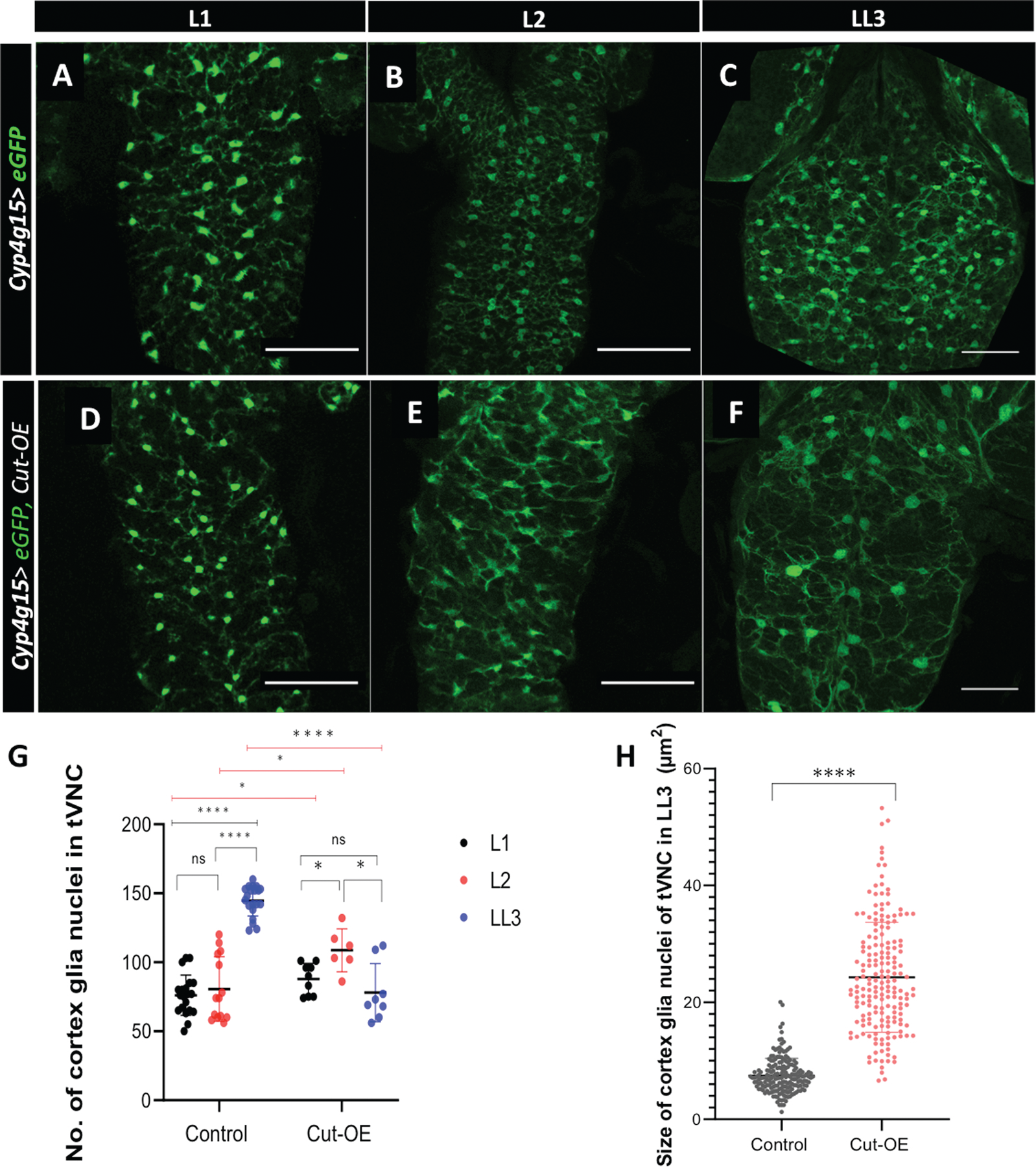
Cut regulates endomitosis of cortex glia. **(A-F)** Confocal projections of the ventral few sections are shown for clarity. Compare the pattern of cortex glia distribution from L1-LL3 in tVNC of control (*Cyp4g15>eGFP/+*)*, (***A-C)** with Cut overexpressing (*Cyp4g15> eGFP, Cut-OE*), **(D-F)**. L1 and L2 control tVNC **(A, B),** the nuclei are arranged more or less in a vertical line pattern (nuclei number drastically increased in LL3 with brighter cell bodies. The distribution of cortex glia in L1 of Cut overexpression is more or less similar to control **(compare A with D),** whereas in L2, it gets a bit distorted (compare **B** with **E**). A striking difference is seen in Cut overexpressing LL3 **(F)** from LL3 control **(C).** The pattern of GFP-positive cortex glia upon Cut overexpression appears more like L1/L2 of controls (compare **D** with **A, B**). No increase in the GFP+ nuclei is observed in the Cut overexpressing tVNC (compare **F** with **C**). **(G)** Further, quantification of cortex glia nuclei from L1-LL3 in control and Cut-OE clearly shows that in Cut overexpression, the nuclei number does not increase from L2 to LL3 (n>5). **(H)** size of the cortex glia nuclei (Repo+, GFP+) was measured by counting 20 nuclei/tVNC (n=5). Statistical evaluation of significance**** if p<0.0001,*** if p<0.001,** if p<0.01 and * if p<0.05 based on unpaired t-test using GraphPad Prism 9 software. Scale bar: 50µm in A-F.

Since we see a similar defect in Cut knockdown where the nuclei number does not increase we set out to check the DNA content upon Cut overexpression. We measured the DNA content through the integrated density of DAPI of each cortex glia nuclei labeled with GFP and Repo (DAPI+, GFP+, Repo+) through ImageJ. Strikingly, the DNA content in each nucleus showed a 2-3fold increase upon Cut overexpression when compared with control (Fig.6 compare C with F, graph J). Thus, Cut overexpressing cortex glia continue to synthesize the DNA which confirms that Cut is necessary and sufficient to increase the DNA content.

**Figure 6.**
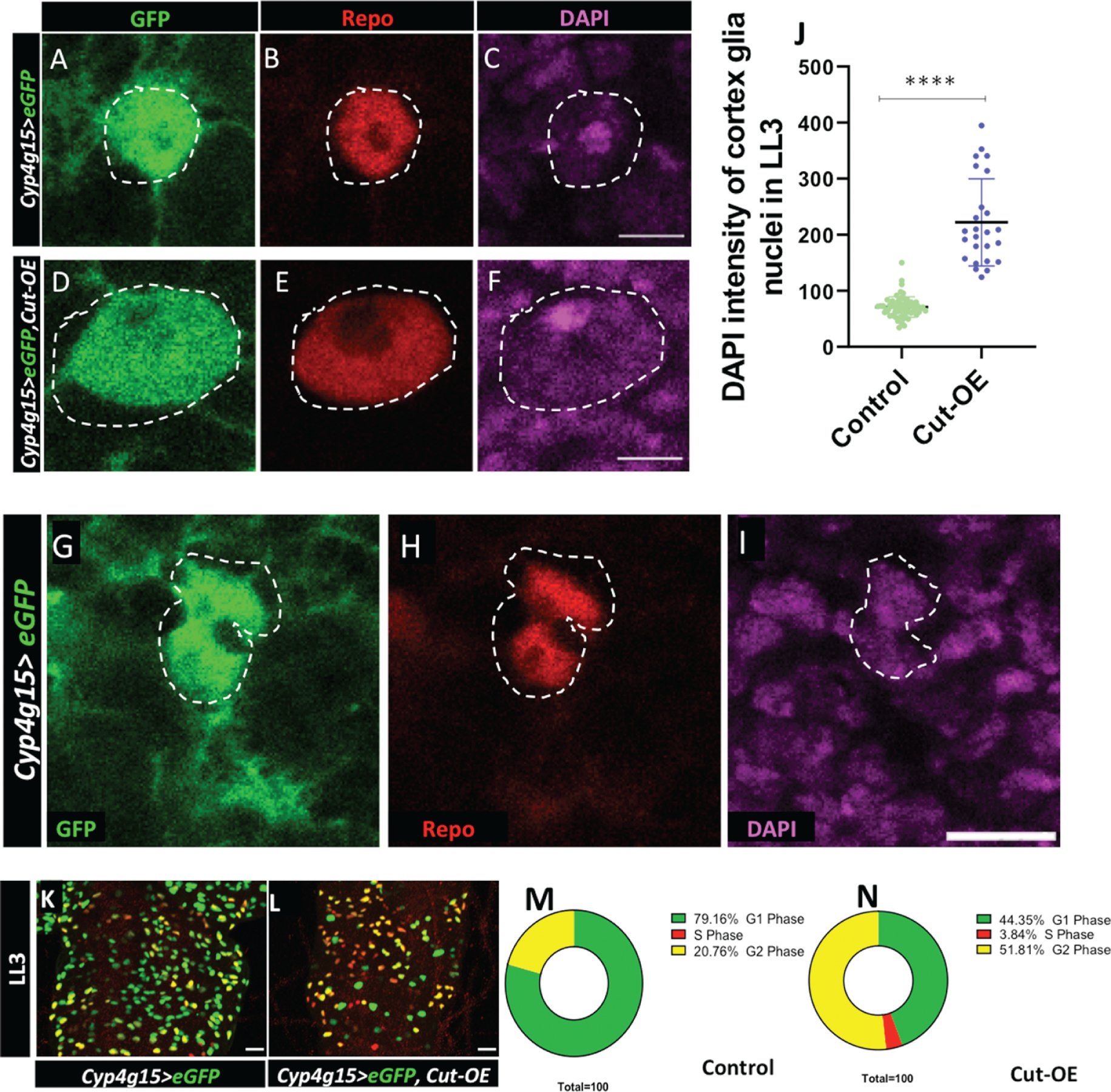
Constitutive activation of Cut increases DNA content in cortex glia. **(A-I)** cortex glia nuclei of LL3 are marked with GFP (green), Repo (Red), and DAPI (magenta). **(A-C)** control cortex glia (*Cyp4g15>eGFP/+),* nuclei of LL3 tVNC **(D-F**) overexpression of Cut (*Cyp4g15>eGFP, Cut-OE*) results in a significant increase in the size of cortex glia nuclei (compare control **A-C** with **D-F** Cut-OE). **(G-I)** control cortex glia appears to be seperating during endomitosis. **(J)** DNA quantification by the integrated density of DAPI in each cortex glia nuclei (more than 20 nuclei counted per CNS (n=3) showing a significant increase in DNA content upon Cut overexpression. **(K-N)** cell cycle stages of cortex glia nuclei in LL3 detected with Fly-FUCCI in **(K)** control (*Cyp4g15>eGFP/+)* and **(L)** (*Cyp4g15>eGFP, Cut-OE*). **(K-N**) fluorescence representing G1 (green), S (red) and G2/M(yellow). **(M-N)** in control count of Nuclei in different cell cycle stages shows that the majority (79.16%) of nuclei are in G1 and rest (20.76%) are in G2/M whereas large number of Cut overexpression nuclei are still in S phase (3.83 %) and also in G2/M phase (51.81%). Scale bar 5µm in **A-I** and 20µm in **K-M**. Statistical evaluation of significance **** p<0.0001, ^ns^p<0.183 based on unpaired t-test.

To get more details about the cell cycle we used the Fly-FUCCI tool to discriminate between G1, S, and G2/M phases of the cell cycle in Cut overexpressing cortex glia. It utilizes fluorescently labeled degrons derived from Cyclin B and E2F1 proteins, which are degraded by APC/C and CRL4/CDT2 proteins from mid-mitosis to the onset of the S phase, respectively (Zielke et al. 2014). At ALH 96hr, most (79.16%) cortex glia in control appears in the G1 phase (Fig.6 K, M). While upon Cut overexpression, a significantly high number of nuclei were still in S (3.83%) and G2/M phase (51.81%) (Fig.6 L, N). Together, it indicates that a large fraction of nuclei in Cut overexpressing glia continue the cell cycle instead of resting at G1, unlike the control (compare Fig. 6 N with M). Therefore, we propose that the homeodomain protein Cut is required for the cortex glia nuclei to increase the DNA content. Probably, the nuclei divide when Cut levels go down, and the cortex glia undergoes endomitosis. The continued high Cut levels lead to endoreplicated nuclei remaining as a single nucleus in a cell.

## Discussion

In this study, we report a novel role of the homeodomain transcription factor Cut in the growth and development of cortex glia. Cortex glia undergo endomitosis and also form a complex reticular network around the neural cells. We observed that Cut regulates growth and DNA content, and loss of Cut interferes with the growth of cell processes and timely increase of DNA content, thus limiting cortex glia to enter endomitosis. On the other hand, continued high Cut levels allow cortex glia to increase their DNA content multifold. We also discovered that despite the increase in DNA content, the constitutively high Cut levels interfere with endomitosis by not allowing these nuclei to separate. We have identified Cut as a novel player in autonomously regulating the dynamic growth of cortex glia around neural cells by controlling cellular morphogenesis and endomitosis.

### Cut a novel regulator of cortex glia morphogenesis

The transcription factor Cut is known to play several roles in the context of cell type and tissue-dependent manner. Acting as a selector gene, it regulates the identity of cells and their growth during development (Krupp et al. 2005). The differential level of Cut and its mammalian homolog, CUX1/2, controls the complexity of dendritic branching, the number of dendritic spines, and tracheal development (Grueber et al. 2003; Cubelos et al. 2010; Pitsouli and Perrimon 2013). Similarly, Cut instructs the growth of cellular processes of the wrapping glia in the peripheral nervous system and acts downstream of FGF signaling (Bauke et al. 2015). In the case of cortex glia, we see that Cut activation enhances the overall growth and thickness of main cortex glia extensions but at the cost of side branching. Our data adds that Cut controls the growth of cortex glia and endomitosis.

It is significant that when cortex glia are ablated in a restricted area, the neighboring cortex glia extends their processes and fills the gaps (Jaeda C. Coutinho-Budd et al. 2017; Hirase et al. 2022). However, Cut defective cortex glia does not show such compensatory growth, resulting in visible wide gaps in the glial trophospongium in the larval CNS. Thus, we conclude that Cut is required and sufficient for the growth of cortex glia processes. However, the signaling pathways and the domain through which Cut functions await further investigation.

Cortex glia processes start growing around the NSCs and neurons of the larval nervous system after receiving the nutritional signals via activation of PI3K/Akt signaling (Spé Der and Brand 2018; Yuan, Conor W. Sipe, et al. 2020). Since Cut defective cortex glia show severely hampered growth of cytoplasmic extensions even as early as L2, we think that Cut may function within the PI3K/Akt signaling pathway in glial cells in a manner analogous to what has been observed in pancreatic cancer cells (Ripka et al. 2010). Other known regulators of cortex glia growth are FGF signaling, components of the membrane fusion machinery, and secreted neurotrophin Spätzle 3 (Read et al. 2009; Witte et al. 2009; Reddy B. V. V. G. and Irvine Kenneth 2011; Avet-Rochex et al. 2012; Jaeda C. Coutinho-Budd et al. 2017; Renee D. Read 2018; Spé Der and Brand 2018; Dong et al. 2020). However, the broad phenotype of Cut activation differs from FGF activation in the cortex glia. Ectopic activation of FGF signaling induces cortex glia overgrowth with a robust increase in nuclei number and overall cell surface area (Avet-Rochex et al. 2012). We found that overexpression of Cut does not enhance the nuclei number of cortex glia. Instead, it resulted in giant nuclei with higher DNA content. Furthermore, overexpression of FGF receptor htlACT in cortex glia makes a heavy network of finer cortex glia extensions (Avet-Rochex et al. 2012). In contrast, Cut overexpression cause a widespread reduction in the finer extensions. Thus, the cytoplasmic growth of fine extension in overexpression of Cut vs. FGF signaling in cortex glia also has different outcomes and may act through distinct mechanisms.

### Regulation of ploidy is essential in the glial cells to control their growth

The size of a cell is important and plays a crucial role during development. Several differentiated cells undergo ploidy changes to increase their size or metabolic output. For example, germline nurse cells, salivary glands, fat bodies, gut, and trachea show varying levels of endopolyploidy (Lilly and Spradling 1996). Interestingly, neurons and glia in invertebrate and vertebrate adult brains compensate for aging-related cell loss or other damages by undergoing polyploidy (Losick et al. 2013; Tamori and Deng 2013; Losick et al. 2016; Cohen et al. 2018; Nandakumar et al. 2020). On the other hand, endomitosis is also a way to increase the cell size through which cells undergo incomplete mitosis and increase the number of nuclei per cell. A few examples of the same are known in *Drosophila* and mammalian systems (Britton and Edgar 1998; Edgar and Orr-Weaver 2001; Taniguchi and Kokuryo 2012; Rios et al. 2016; von Stetina et al. 2018; Windmueller et al. 2020). Several reports from independent labs strongly indicate that cortex glia increase their nuclei number by undergoing endomitosis (Jaeda C Coutinho-Budd et al. 2017; Yuan, Conor W Sipe, et al. 2020; Rujano et al. 2022). A recent study shows that cortex glia make syncytia and complex networks by fusion and undergoing endocycle and endomitosis (Rujano et al. 2022). Cut regulates the growth of cortex glia by regulating the complexity of cytoplasmic processes and interferes with endomitosis. The cortex glia having Cut knockdown shows a clear defect in DNA increase and remains stuck in the earlier development stage. Their nuclei do not divide or enter the endomitosis. Interestingly overexpression of Cut clearly shows an increase in DNA content yet these nuclei do not divide. Thus, in cortex glia, over-expression of Cut does not show the routine increase of nuclei number from the L2 to LL3 stage as seen in control tVNC; instead, it increases the ploidy in nuclei. In the process of *Drosophila* oogenesis, follicular cells undergo endocycle to increase their DNA content. To ensure proper development, the downregulation of Cut is necessary to switch from mitosis to endocycle (Sun and Deng 2005). Interestingly, the role of Cut in cortex glia cells is different, as high levels of Cut promote DNA increase per nuclei. This suggests that Cut may have diverse and tissue-specific effects on the cell cycle.

Aneuploidy is one of the most common phenomena seen in several tumors (Rajagopalan and Lengauer 2004), and Cut overexpression is also (Sansregret and Nepveu 2008) reported in diverse classes of cancer (Sansregret, and Nepveu 2008, Ramdzan and Nepveu 2014). There are several tumors, including glioma, where CUX1 overexpression has been reported and considered among driver mutation (Sansregret and Nepveu 2008; Kaur et al. 2018; Feng et al. 2021; Griesmann et al. 2021; Xu et al. 2021). A *in vitro* study shows that mammalian CUX1 has the potential to cause chromosomal instability (Sansregret et al. 2011). Our findings highlight that Cut enhances the DNA content of cortex glia. It would be interesting to see if the genetic combination of Cut with any other tumor driver identified in glioma could make the nuclei unstable and lead to cancer in the brain. In conclusion, our finding that Cut regulates cortex glia growth and endomitosis is of significant importance for development and disease prospects.

## Materials and methods

### Fly stocks and genotypes

*Drosophila melanogaster* was reared at 22^0^C, and RNAi crosses were done at 29°C on standard food medium containing sugar, agar, maize powder, and yeast. Appropriate fly crosses were set up following standard methods to obtain progeny of desired genotypes. The following fly stocks are used in the experiments: Oregon R^+^ as wild-type, *Repo-GAL4* (BL 7415), *Cut-RNAi* (BL 33967, strong), *Cut-RNAi* (BL29625, weak), *UAS-Cut* (II, referred in text as Cut-OE) (Norbert Perrimon, Harvard Medical School, MA, USA), *Cyp4g15-GAL4 (*39103), *UAS-eGFP*(II) (BL-5431), *R54H02-GAL4* (BL-45784), *R46H12-GAL4* (BL-50285), *NP577-GAL4* (Awasaki et al. 2008), *mCD8RFP* (BL-27398), Fly-FUCCI (BL-55121).The following genotype combinations are generated in the lab for the study: *UAS-eGFP*; *Cyp4g15-GAL4*, *UAS-FUCCI;Cyp4g15-GAL4*, *mCD8RFP; Cyp4g15-GAL4, UAS-eGFP; Cut-RNAi*.

### Survival assay

We used three strains (*Oregon R^+^, Cyp4g15-GAL4, and Cut-RNAi*) reared on standard agar-cornmeal-sugar-yeast food. We crossed 60 male and 90 virgin fruit flies. The *Cyp4g15-GAL4* strain was used as males and the *Oregon R^+^* and *Cut-RNAi* strains were used as virgin females. The flies were isolated, fed, and reared for two days before being transferred to a food bottle for mating. We placed 20 males and 30 females in each replicate bottle and allowed them to mate for a day. After mating, the flies were transferred to a cage with an agar plate where they laid eggs for 12hr. The agar plates were changed daily. To analyze survival at the pupariation stage and eclosion, we collected synchronized 1st instar larvae and transferred them to food vials. We examined 11 replicates of 100 larvae each for the control and *Cut-RNAi* groups. The percentage of pupation and eclosion was calculated as described earlier (Kharat et al. 2020). For developmental delay analysis, we examined only two replicates of 100 larvae for the control and *Cut-RNAi* groups. At least three replicates of 100 larvae each were examined in each group for the survival assay, with one to two-day-old imagoes placed in food vials. The recorded data were analyzed using GraphPad Prism 9 software for statistical analysis.

### Immunostaining, confocal microscopy, and documentation

Larvae of the required age from F1 progeny of respective crosses were selected, and CNS were dissected in Phosphate Buffer (PBS 1X containing-NaCl, KCL, NA2HPO4, KH2PO4, pH-7.4), fixed in 4% Paraformaldehyde for 30 min., rinsed in 0.1% PBST (1XPBS,0.1%TritonX-100), and incubated in blocking solution (0.1% TritonX-100, 0.1%BSA, 10%FCS, 0.1% deoxycholate, 0.02% thiomersal) for 30min., at room temperature. Samples were incubated in the required primary antibody at four-degree centigrade overnight. Antibodies used are mouse anti-Cut (1:10, Developmental Studies Hybridoma Bank, 2B10), mouse, Anti-Repo (1:50, Developmental Studies Hybridoma Bank, 8D12), rabbit anti-GFP antibody (1:200, Invitrogen, A-11122). For rat anti-Dpn antibody (1:150, ab195173, Abcam), the samples were kept for 2 consecutive overnights for better signal. The following day, samples were rinsed thrice in 0.1% PBST and incubated with appropriate secondary antibody at 1:200 dilution overnight or two hours at room temperature. Secondary antibodies used are donkey anti-rat Alexa 488 (Invitrogen, A-21208), Goat anti-Mouse Alexa Fluor 568 (cat. No. A11004), Chicken Anti-Rabbit Alexa Fluor 488 (Invitrogen, A-21441). Following incubation with secondary antibodies, the samples were washed thrice in 0.1%PBST, counterstained with DAPI (1µg/ml, Invitrogen, D1306) at 4°C overnight whenever needed. After the final washes in 0.1%PBST, the samples were mounted in DABCO (Sigma, D27802) for further analysis.

Images acquired at the following confocal microscopes Zeiss LSM-510 Meta at the Department of Zoology, Zeiss LSM-510 at ISLS BHU, and Leica SP8 STED facility at CDC, BHU. All the images were quantified using Fiji/ImageJ software (NIH, USA). Images were assembled using adobe photoshop and MS PowerPoint softwares.

### Image analysis using Fiji/ImageJ application

Confocal sections of cortex glia of tVNC were analyzed using Fiji/ImageJ software (NIH, USA) to measure number, area, and volume. For integrated intensity measurements in a region of interest (ROI) freehand selection tool was used to mark the area manually.

The number of Glia nuclei and NSCs were counted manually based on GAL4-driven GFP and Repo staining using the multipoint selection tool. To measure the nuclei size of cortex glia (GFP+, Repo+) was used to outline the area. The area outline was done using the freehand selection tool around Repo staining, followed by selecting an analyse-measure option. Chamber size was measured by outlining the cortex glia processes that are coming out from the cell body around one nucleus and forming a chamber around progeny using a freehand selection tool, size of the outlined area (µm^2^) was measured using the analyze-measure option. For volume analysis of cortex glia, one specific threshold was chosen that fits best for the actual staining and was kept constant throughout the analysis. The thresholding procedure is used in image processing to select pixels of interest based on the GFP intensity of the pixel values. Thereafter measure stack plugin was used to find the fluorescent area (GFP) of each cortex glia section of tVNC through the analyze-measure option where Area, mean gray value, stack position, and limit to threshold checkboxes were selected. The area obtained was thus multiplied by the number of stack intervals (=2) to find out the volume of cortex glia. Finally, the area covered was estimated by manually outlining the area occupied by tVNC.

The density of cortex glia processes was calculated through the plot profile and the surface profile options. The tools measure the signal intensities of cortex glia processes along the selected lines. The gray values indicate the intensities of cortex glia along a line, and the number of peaks denotes the number of cortex glia extensions.

Integrated density for DAPI in the cortex glia nuclei was measured by manually outlining each GFP+, Repo+ DAPI+ nuclei, the area outline was done using the freehand selection tool around Repo staining. The integrated density of DAPI was measured in each section of tVNC using the analyze-measure option. The confocal scanning parameters for each experimental setup were kept constant for all intensity quantifications. Controls were analyzed in parallel to each experimental set.

Statistical analysis was done using GraphPad Prism 9 software where a two-tailed unpaired t-test was used to evaluate the statistical significance between the mean values of the two groups, using asterisk as follows: n with p < 0.05 (**** if p<0.0001,*** if p<0.001,** if p<0.01 and * if p<0.05) considered statistically significant and if p>0.05 considered non-significant

## Data Availability

Fly lines are obtained from Bloomington Stock Center, USA, and genetic combinations created are available upon request. The authors affirm that all other data necessary for confirming the conclusions of the article are present within the article, figures, and tables. Supplemental material available at GENETICS online.

## Acknowledgment

We are grateful to Profs. S C Lakhotia, P. Sinha, A. Mukherjee, Drs. A. Verma, B. C. Mondal, and G Pandey, for critically reviewing the manuscript, insightful comments, and valuable suggestions throughout. We also acknowledge Prof A. Nepveu, Prof. M. Freeman, and Dr. Jaeda C. Coutinho-Budd, for their communication, which helped us develop and refine our hypothesis. We also thank the N. Perrimon, T. Awasaki, and Bloomington Drosophila Stock Center (BDSC) for providing fly stocks. The national facility of Zeiss 510 Meta confocal microscopy at the Department of Zoology, and ISLS BHU, Leica SP8 STED confocal microscopy facility at CDC, BHU are also acknowledged. Finally, we thank the Cytogenetics Laboratory, Department of Zoology, for all the instrumentation facilities.

## Author’s contribution

All authors contributed to the article. VY performed the animal experiments, image acquisition, design of the quantification workflow, statistical analysis, and made the figure panels under the supervision of RA. Part of the quantification of Fig. 1k,3J, and Fig.S3A-B. is done by RKM under VY and RA supervision. PD has re-analyzed the quantification of Fig. 4G, 6J under the supervision of VY and RA.

**Figure.S1 Cut expresses in glial subtypes and maintains niche around neural stem cells**

**(A-B)** compare the membrane network in LL3 **(A)** control *(Cyp4g15>eGFP/+),* **(B)** Cut knockdown weak RNAi line *(Cyp4g15>eGFP; Cut-RNAi),* the cortex glia networks get disrupted even with weak RNAi line (BL#29625). **(C)** Quantification of NSCs upon Cut-knockdown (strong RNAi line BL # 33967) with Repo-GAL4 and several cortex glia-specific drivers (*Cyp4g15-GAL4, R54H02-GAL4, R46H12-GAL4, NP577-GAL4*). **(D-I)** Single confocal sections showing Cut expression (red) in **(D-F)** pan glia marked with *Repo>eGFP*, and in **(G-I)** cortex glia marked with *Cyp4g15>eGFP*. Statistical evaluation of significance. **** p<0.0001, ** 0.0041, ** 0.0012 based on unpaired t-test.

**Figure S2: Cut knockdown in the cortex glia leads to loss of cortex glia processes and nuclei in the whole CNS**

**(A-A’)** Control *(Cyp4g15>eGFP/+)* showing fine honeycomb structure of cortex glia processes, **(B-B’)** Cut defective cortex glia (*Cyp4g15>eGFP, Cut-RNAi*) with minimum processes, **(A’, B**’ are the merged projections with Repo to show glia). **(C, D)** represent the surface plot of cortex glia in tVNC to compare the intensity and extensions of **(C)** control with **(D)** Cut defective cortex glia. **(E-H’’)** loss of Cut in cortex glia affects their process in the abdominal (control **G-G”** compare with **H-H”)** and optical lobes (control **E-E”** compare with **F-F”). (I, I’)** Late 16^th^ stage embryo (*Cyp4g15>mCD8RFP)* showing that GAL4 starts expressing from embryonic life onwards, cortex glia (red), Repo(green). **(J)** quantification of cortex glia nuclei in the abdominal region of VNC during different larval time points (ALH 24hr and LL3), in control *(Cyp4g15> eGFP),* and upon Cut knockdown *(Cyp4g15> eGFP;Cut-RNAi).* Note in LL3 the nuclei number does not significantly increase upon Cut knockdown as seen in control. Statistical evaluation of significance *p= 0.0271, ****p<0.0001, based on unpaired t-test using GraphPad prism 9 software. Scale bar: 50µm in A-B’ and 20µm in I, I’.

**Figure S3: Cut defective Cortex glia is associated with poor survival and developmental delay.**

the genotypes of control (*Cyp4g15>eGFP*) and Cut knockdown (*Cyp4g15>eGFP; Cut-RNAi*) throughout the data.

**(A)** Percentage developmental delay, statistical evaluation of significance *p= 0.0206, using paired t-test and ^ns^p> 0.05 using unpaired t-test, **(B)** Effect of Cut knockdown on the lifespan of *Drosophila* adult flies. Median life shows a significant difference ** indicates P ≤ 0.0056 between the Control & *Cut-RNAi* using log-rank (Mantel–Cox) test.**(C-D’)** projection of ventral sections of tVNC at ALH 72hr in control and Cut knockdown, shows major loss of cortex glia processes and are unable to form fine branches upon Cut knockdown.

## References

1. Abbott NJ. 2005. Dynamics of CNS Barriers: Evolution, Differentiation, and Modulation. Cellular and Molecular Neurobiology 2005 25:1. 25(1):5–23. doi:10.1007/S10571-004-1374-Y. https://link.springer.com/article/10.1007/s10571-004-1374-y.

2. Andres V, Bernardo nadal-ginard, Vijak mahdavi. 1992. Clox, a mammalian homeobox gene related to Drosophila cut, encodesDNA-binding regulatory proteins differentially expressed duringdevelopment. Development .:321–334.

3. Arya R, Gyonjyan S, Harding K, Sarkissian T, Li Y, Zhou L, White K. 2019. A Cut/cohesin axis alters the chromatin landscape to facilitate neuroblast death. Development (Cambridge). 146(9). doi:10.1242/dev.166603.

4. Avet-Rochex A, Kaul AK, Gatt AP, McNeill H, Bateman JM. 2012. Concerted control of gliogenesis by InR/TOR and FGF signalling in the Drosophila post-embryonic brain. Development. 139(15):2763–2772. doi:10.1242/dev.074179.

5. Awasaki T, Lai SL, Ito K, Lee T. 2008. Organization and postembryonic development of glial cells in the adult central brain of Drosophila. Journal of Neuroscience. 28(51):13742–13753. doi:10.1523/JNEUROSCI.4844-08.2008. https://www.jneurosci.org/content/28/51/13742.

6. Barres BA. 2008. The Mystery and Magic of Glia: A Perspective on Their Roles in Health and Disease. Neuron. 60(3):430–440. doi:10.1016/J.NEURON.2008.10.013/ATTACHMENT/A48AE9A5-41E7-4DAA-B6BB-75687BD12A10/MMC1.PDF. http://www.cell.com/article/S0896627308008866/fulltext.

7. Bauke AC, Sasse S, Matzat T, Klämbt C. 2015. A transcriptional network controlling glial development in the drosophila visual system. Development (Cambridge). 142(12):2184–2193. doi:10.1242/dev.119750.

8. Blochlinger K, Bodmer R, Jack J, Jan LY, Jan YN. 1988. Primary structure and expression of a product from cut, a locus involved in specifying sensory organ identity in Drosophila. Nature. 333(6174):629–635. doi:10.1038/333629A0. https://pubmed.ncbi.nlm.nih.gov/2897632/.

9. Bodmer R, Barbel S, Sheperd S, Jack JW, Jan LY, Jan YN. 1987. Transformation of sensory organs by Mutations of the cut locus of D. melanogaster. Cell. 51(2):293– 307. doi:10.1016/0092-8674(87)90156-5.

10. Booth GE, Kinrade EF v, Hidalgo A. 2000. Glia maintain follower neuron survival during Drosophila CNS development. Development. 127(2):237–244. doi:10.1242/DEV.127.2.237. https://journals.biologists.com/dev/article/127/2/237/40942/Glia-maintain-follower-neuron-survival-during.

11. Britton JS, Edgar BA. 1998. Environmental control of the cell cycle in Drosophila: Nutrition activates mitotic and endoreplicative cells by distinct mechanisms. Development. 125(11):2149–2158.

12. Cadieux C, Kedinger V, Yao L, Vadnais C, Drossos M, Paquet M, Nepveu A. 2009. Mouse mammary tumor virus p75 and p110 CUX1 transgenic mice develop mammary tumors of various histologic types. Cancer Res. 69(18):7188–7197. doi:10.1158/0008-5472.CAN-08-4899. https://pubmed.ncbi.nlm.nih.gov/19738070/.

13. Cohen E, Allen SR, Sawyer JK, Fox DT. 2018. Fizzy-Related dictates A cell cycle switch during organ repair and tissue growth responses in the Drosophila hindgut. Elife. 7. doi:10.7554/eLife.38327. https://elifesciences.org/articles/38327.

14. Coqueret O, Bérubé G, Nepveu A. 1998. The mammalian Cut homeodomain protein functions as a cell-cycle-dependent transcriptional repressor which downmodulates p21(WAF1/CIP1/SDI1) in S phase. EMBO Journal. 17(16):4680– 4694. doi:10.1093/emboj/17.16.4680. https://pubmed.ncbi.nlm.nih.gov/9707427/.

15. Coutinho-Budd Jaeda C., Sheehan AE, Freeman MR. 2017. The secreted neurotrophin spätzle 3 promotes glial morphogenesis and supports neuronal survival and function. Genes Dev. 31(20):2023–2038. doi:10.1101/gad.305888.117.

16. Cubelos B, Sebastián-Serrano A, Beccari L, Calcagnotto ME, Cisneros E, Kim S, Dopazo A, Alvarez-Dolado M, Redondo JM, Bovolenta P, et al. 2010. Cux1 and Cux2 regulate dendritic branching, spine morphology, and synapses of the upper layer neurons of the cortex. Neuron. 66(4):523–535. doi:10.1016/j.neuron.2010.04.038.

17. Dong Q, Zavortink M, Froldi F, Golenkina S, Lam T, Cheng LY. 2021. Glial Hedgehog signalling and lipid metabolism regulate neural stem cell proliferation in Drosophila . EMBO Rep. 22(5). doi:10.15252/embr.202052130.

18. Dumstrei K, Wang F, Hartenstein V. 2003a. Role of DE-Cadherin in Neuroblast Proliferation, Neural Morphogenesis, and Axon Tract Formation in Drosophila Larval Brain Development.

19. Edgar BA, Orr-Weaver TL. 2001. Endoreplication Cell Cycles: Review More for Less.

20. Feng F, Zhao Z, Zhou Y, Cheng Y, Wu X, Heng X. 2021. CUX1 Facilitates the Development of Oncogenic Properties Via Activating Wnt/β-Catenin Signaling Pathway in Glioma. Front Mol Biosci. 8:764. doi:10.3389/fmolb.2021.705008.

21. Freeman MR, Doe CQ. 2001. Asymmetric Prospero localization is required to generate mixed neuronal/glial lineages in the Drosophila CNS. Development. 128(20):4103–4112. doi:10.1242/dev.128.20.4103. https://pubmed.ncbi.nlm.nih.gov/11641232/.

22. Gonzalez-Gutierrez A, Ibacache A, Esparza A, Barros LF, Sierralta J. 2019 Apr. Monocarboxylate Transport in Drosophila Larval Brain during Low and High Neuronal Activity. bioRxiv.:610196. doi:10.1101/610196.

23. Griesmann H, Mühl S, Riedel J, Theuerkorn K, Sipos B, Esposito I, Vanden Heuvel GB, Michl P. 2021. CUX1 Enhances Pancreatic Cancer Formation by Synergizing with KRAS and Inducing MEK/ERK-Dependent Proliferation. Cancers (Basel). 13:2462. doi:10.3390/cancers13102462.

24. Grueber WB, Jan LY, Nung Jan Y. 2003. Different Levels of the Homeodomain Protein Cut Regulate Distinct Dendrite Branching Patterns of Drosophila Multidendritic Neurons.

25. vanden Heuvel GB, Brantley JG, Alcalay NI, Sharma M, Kemeny G, Warolin J, Ledford AW, Pinson DM. 2005. Hepatomegaly in Transgenic Mice Expressing the Homeobox Gene Cux-1. Mol Carcinog. 43(1):18. doi:10.1002/MC.20091. /pmc/articles/PMC4441415/.

26. Hirase H, López-Hidalgo M, Klämbt C, Coutinho-Budd J, Salazar G, Ross G, Maserejian AE. 2022. Quantifying Glial-Glial Tiling Using Automated Image Analysis in Drosophila. Automated Image Analysis in Drosophila Front Cell Neurosci. 16:826483. doi:10.3389/fncel.2022.826483. www.frontiersin.org.

27. Ito K, Urban J, Technau GM. 1995. Distribution, classification, and development of Drosophila glial cells in the late embryonic and early larval ventral nerve cord. Roux’s Archives of Developmental Biology. 204(5):284–307. doi:10.1007/BF02179499.

28. Kaur S, Ramdzan ZM, Guiot MC, Li L, Leduy L, Ramotar D, Sabri S, Abdulkarim B, Nepveu A. 2018. CUX1 stimulates APE1 enzymatic activity and increases the resistance of glioblastoma cells to the mono-alkylating agent temozolomide. Neuro Oncol. 20(4):484–493. doi:10.1093/neuonc/nox178. https://academic.oup.com/neuro-oncology/article/20/4/484/4237503.

29. Kharat P, Sarkar P, Mouliganesh S, Tiwary V, Priya VBR, Sree NY, Annapoorna HV, Saikia DK, Mahanta K, Thirumurugan K. 2020. Ellagic acid prolongs the lifespan of Drosophila melanogaster. Geroscience. 42(1):271–285. doi:10.1007/s11357-019-00135-6.

30. Krupp JJ, Yaich LE, Wessells RJ, Bodmer R. 2005. Identification of Genetic Loci That Interact With cut During Drosophila Wing-Margin Development. doi:10.1534/genetics.105.043125. https://academic.oup.com/genetics/article/170/4/1775/6060441.

31. Ledford AW, Brantley JG, Kemeny G, Foreman TL, Quaggin SE, Igarashi P, Oberhaus SM, Rodova M, Calvet JP, vanden Heuvel GB. 2002. Deregulated expression of the homeobox gene Cux-1 in transgenic mice results in downregulation of p27(kip1) expression during nephrogenesis, glomerular abnormalities, and multiorgan hyperplasia. Dev Biol. 245(1):157–171. doi:10.1006/DBIO.2002.0636. https://pubmed.ncbi.nlm.nih.gov/11969263/.

32. Lilly MA, Spradling AC. 1996. The Drosophila endocycle is controlled by Cyclin E and lacks a checkpoint ensuring S-phase completion.

33. Liu S, Jack J. 1992. Regulatory interactions and role in cell type specification of the Malpighian tubules by the cut, Krüppel, and caudal genes of Drosophila. Dev Biol. 150(1):133–143. doi:10.1016/0012-1606(92)90013-7.

34. Liu S, McLeod E, Jack J. 1991. Four Distinct Regulatory Regions of the Cut Locus and Their Effect on Cell Type Specification in Drosophila. Genetics. 127(1):151. doi:10.1093/GENETICS/127.1.151. /pmc/articles/PMC1204300/?report=abstract.

35. Losick VP, Fox DT, Spradling AC. 2013. Polyploidization and cell fusion contribute to wound healing in the adult Drosophila epithelium. Current Biology. 23(22):2224–2232. doi:10.1016/j.cub.2013.09.029.

36. Losick VP, Jun AS, Spradling AC, Dr M. 2016. Wound-Induced Polyploidization: Regulation by Hippo and JNK Signaling and Conservation in Mammals. doi:10.1371/journal.pone.0151251.

37. Ludlow C, Choy R, Blochlinger K. 1996. Functional Analysis of Drosophila and Mammalian Cut Proteins in Flies. Dev Biol. 178(1):149–159. doi:10.1006/DBIO.1996.0205.

38. Michl P, Ramjaun AR, Pardo OE, Warne PH, Wagner M, Poulsom R, D’Arrigo C, Ryder K, Menke A, Gress T, et al. 2005. CUTL1 is a target of TGFβ signaling that enhances cancer cell motility and invasiveness. Cancer Cell. 7(6):521–532. doi:10.1016/j.ccr.2005.05.018. https://pubmed.ncbi.nlm.nih.gov/15950902/.

39. Nandakumar S, Grushko O, Buttitta LA. 2020. Polyploidy in the adult drosophila brain. Elife. 9:1–25. doi:10.7554/ELIFE.54385.

40. Nepveu A. 2001. Role of the multifunctional CDP/Cut/Cux homeodomain transcription factor in regulating differentiation, cell growth and development. Gene. 270(1–2):1–15. doi:10.1016/S0378-1119(01)00485-1.

41. Neufeld EJ, Skalnik DG, Lievens PMJ, Orkin SH. 1992. Human CCAAT displacement protein is homologous to the Drosophila homeoprotein, cut. Nat Genet. 1(1):50–55. doi:10.1038/NG0492-50.

42. Oland LA, Tolbert LP. 2003. KEY INTERACTIONS BETWEEN NEURONS AND GLIAL CELLS DURING NEURAL DEVELOPMENT IN INSECTS. Annu Rev Entomol. 48:89–110. doi:10.1146/annurev.ento.48.091801.112654. www.annualreviews.org.

43. Pereanu W, Shy D, Hartenstein V. 2005a. Morphogenesis and proliferation of the larval brain glia in Drosophila. Dev Biol. 283(1):191–203. doi:10.1016/j.ydbio.2005.04.024.

44. Pitsouli C, Perrimon N. 2013. The homeobox transcription factor cut coordinates patterning and growth during drosophila airway remodeling. Sci Signal. 6(263):ra12. doi:10.1126/scisignal.2003424. https://www.science.org/doi/10.1126/scisignal.2003424.

45. Quaggin SE, vanden Heuvel GB, Golden K, Bodmer R, Igarashi P. 1996. Primary structure, neural-specific expression, and chromosomal localization of Cux-2, a second murine homeobox gene related to Drosophila cut. Journal of Biological Chemistry. 271(37):22624–22634. doi:10.1074/JBC.271.37.22624.

46. Rajagopalan H, Lengauer C. 2004. Aneuploidy and cancer. Nature. 432(7015):338–341. doi:10.1038/NATURE03099. https://pubmed.ncbi.nlm.nih.gov/15549096/.

47. Ramdzan ZM, Nepveu A. 2014. CUX1, a haploinsufficient tumour suppressor gene overexpressed in advanced cancers. Nat Rev Cancer. 14(10):673–682. doi:10.1038/nrc3805.

48. Read Renee D. 2018. Pvr receptor tyrosine kinase signaling promotes post-embryonic morphogenesis, and survival of glia and neural progenitor cells in drosophila. Development (Cambridge). 145(23):dev164285. doi:10.1242/dev.164285. http://www.ncbi.nlm.nih.gov/pubmed/30327326.

49. Read RD, Cavenee WK, Furnari FB, Thomas JB. 2009. A Drosophila model for EGFR-Ras and PI3K-dependent human glioma. PLoS Genet. 5(2). doi:10.1371/journal.pgen.1000374.

50. Reddy B. V. V. G., Irvine Kenneth. 2011. Regulation of Drosophilaglial cell proliferation by Merlin-Hippo signaling. Development. 138:5201–5212. doi:10.1242/dev.069385.

51. Rios AC, Fu NY, Jamieson PR, Pal B, Whitehead L, Nicholas KR, Lindeman GJ, Visvader JE. 2016. Essential role for a novel population of binucleated mammary epithelial cells in lactation. Nat Commun. 7. doi:10.1038/NCOMMS11400. /pmc/articles/PMC4844753/.

52. Rujano MA, Briand D, Ðelić B, Marc J, Spéder P. 2022a. An interplay between cellular growth and atypical fusion defines morphogenesis of a modular glial niche in Drosophila. Nat Commun. 13(1):1–25. doi:10.1038/s41467-022-32685-3. https://www.nature.com/articles/s41467-022-32685-3.

53. Sansregret L, Nepveu A. 2008a. The multiple roles of CUX1: Insights from mouse models and cell-based assays. Gene. 412(1–2):84–94. doi:10.1016/J.GENE.2008.01.017.

54. Sansregret L, Vadnais C, Livingstone J, Kwiatkowski N, Awan A, Cadieux C, Leduy L, Hallett MT, Nepveu A. 2011. Cut homeobox 1 causes chromosomal instability by promoting bipolar division after cytokinesis failure. Proc Natl Acad Sci U S A. 108(5):1949–1954. doi:10.1073/pnas.1008403108.

55. Spé Der P, Brand AH. 2018. Systemic and local cues drive neural stem cell niche remodelling during neurogenesis in Drosophila. Elife. doi:10.7554/eLife.30413.001. https://doi.org/10.7554/eLife.30413.001.

56. von Stetina JR, Frawley LE, Unhavaithaya Y, Orr-Weaver TL. 2018. Variant cell cycles regulated by Notch signaling control cell size and ensure a functional blood-brain barrier. Development (Cambridge). 145(3). doi:10.1242/DEV.157115/19236. https://journals.biologists.com/dev/article/145/3/dev157115/19236/Variant-cell-cycles-regulated-by-Notch-signaling.

57. Stork T, Bernardos R, Freeman MR. 2012. Analysis of glial cell development and function in Drosophila. Cold Spring Harb Protoc. 7(1):1–17. doi:10.1101/pdb.top067587.

58. Sun J, Deng WM. 2005. Notch-dependent downregulation of the homeodomain gene cut is required for the mitotic cycle/endocycle switch and cell differentiation in Drosophila follicle cells. Development. 132(19):4299–4308. doi:10.1242/dev.02015. https://pubmed.ncbi.nlm.nih.gov/16141223/.

59. Tamori Y, Deng WM. 2013. Tissue Repair through Cell Competition and Compensatory Cellular Hypertrophy in Postmitotic Epithelia. Dev Cell. 25(4):350–363. doi:10.1016/j.devcel.2013.04.013.

60. Taniguchi K, Kokuryo A. 2012. Binucleation of Drosophila Adult Male Accessory Gland Cells Increases Plasticity of Organ Size for Effective Reproduction. Biol Syst Open Access. 01(01). doi:10.4172/2329-6577.1000e101.

61. Tavares AT, Tsukui T, Belmonte JCI. 2000. Evidence that members of the Cut/Cux/CDP family may be involved in AER positioning and polarizing activity during chick limb development. Development. 127(23):5133–5144. doi:10.1242/DEV.127.23.5133. https://journals.biologists.com/dev/article/127/23/5133/41115/Evidence-that-members-of-the-Cut-Cux-CDP-family.

62. Trapp BD, Nave KA. 2008. Multiple sclerosis: An immune or neurodegenerative disorder? Annu Rev Neurosci. 31:247–269. doi:10.1146/annurev.neuro.30.051606.094313.

63. Unhavaithaya Y, Orr-Weaver TL. 2012. Polyploidization of glia in neural development links tissue growth to blood–brain barrier integrity. Genes Dev. 26(1):31. doi:10.1101/GAD.177436.111. /pmc/articles/PMC3258963/.

64. Vadnais C, Davoudi S, Afshin M, Harada R, Dudley R, Clermont PL, Drobetsky E, Nepveu A. 2012. CUX1 transcription factor is required for optimal ATM/ATR-mediated responses to DNA damage. Nucleic Acids Res. 40(10):4483–4495. doi:10.1093/nar/gks041.

65. Valarche, J P Tissier-Seta, M R Hirsch SM, C Goridis, J F Brunet. 1993. The mouse homeodomain protein Phox2 regulates N cam promoter activity in concert with Cux/CDP and is a putative determinant of neurotransmitter phenotype. Development.:881–896.

66. Weiss S, Melom JE, Ormerod KG, Zhang Y v, Littleton JT. 2019. Glial Ca2+signaling links endocytosis to K+ buffering around neuronal somas to regulate excitability. Elife. 8. doi:10.7554/eLife.44186. https://elifesciences.org/articles/44186.

67. van Wijnen AJ, van Gurp MF, de Ridder MC, Tufarellit C, Last TJ, Birnbaum M, Vaughan PS, Giordanot A, Krek W, Neufeldt EJ, et al. 1996. CDP/cut is the DNA-binding subunit of histone gene transcription factor HiNF-D: A mechanism for gene regulation at the G1/S phase cell cycle transition point independent of transcription factor E2F (proliferation/gene expression/cyclin-dependent kinase/tumor suppressor). Biochemistry. 93:11516–11521. [accessed 2022 Oct 18]. https://www.pnas.org.

68. Windmueller R, Leach JP, Babu A, Zhou S, Morley MP, Wakabayashi A, Petrenko NB, Viatour P, Morrisey EE. 2020. Direct Comparison of Mononucleated and Binucleated Cardiomyocytes Reveals Molecular Mechanisms Underlying Distinct Proliferative Competencies. Cell Rep. 30(9):3105. doi:10.1016/J.CELREP.2020.02.034. /pmc/articles/PMC7194103/.

69. Witte HT, Jeibmann A, Klämbt C, Paulus W. 2009. Modeling glioma growth and invasion in Drosophila melanogaster. Neoplasia. 11(9):882–888. doi:10.1593/neo.09576.

70. Xiong W-C, Montell C. 1995. Defective Glia Induce Neuronal Apoptosis in the repo Visual System of Drosophila. Neuron. 14:581–590.

71. Xu A, Wang X, Luo J, Zhou M, Yi R, Huang T, Lin J, Wu Z, Xie C, Ding S, et al. 2021. Overexpressed P75CUX1 promotes EMT in glioma infiltration by activating β-catenin. Cell Death Dis. 12(2):1–15. doi:10.1038/s41419-021-03424-1. https://www.nature.com/articles/s41419-021-03424-1.

72. Yoon SO, Chikaraishi DM. 1994. Isolation of two E-box binding factors that interact with the rat tyrosine hydroxylase enhancer. Journal of Biological Chemistry. 269(28):18453–18462. doi:10.1016/s0021-9258(17)32330-x.

73. Yuan X, Sipe Conor W., Suzawa M, Bland ML, Siegrist SE. 2020. Dilp-2-mediated PI3-kinase activation coordinates reactivation of quiescent neuroblasts with growth of their glial stem cell niche. PLoS Biol. 18(5). doi:10.1371/journal.pbio.3000721.

74. Zhai Z, Ha N, Papagiannouli F, Hamacher-Brady A, Brady N, Sorge S, Bezdan D, Lohmann I. 2012. Antagonistic regulation of apoptosis and differentiation by the cut transcription factor represents a tumor-suppressing mechanism in drosophila. PLoS Genet. 8(3):e1002582. doi:10.1371/journal.pgen.1002582. https://journals.plos.org/plosgenetics/article?id=10.1371/journal.pgen.1002582.

75. Zhou B, Zuo YX, Jiang RT. 2019. Astrocyte morphology: Diversity, plasticity, and role in neurological diseases. CNS Neurosci Ther. 25(6):665–673. doi:10.1111/cns.13123.

76. Zielke N, Korzelius J, vanStraaten M, Bender K, Schuhknecht GFP, Dutta D, Xiang J, Edgar BA. 2014. Fly-FUCCI: A Versatile Tool for Studying Cell Proliferation in Complex Tissues. Cell Rep. 7(2):588–598. doi:10.1016/j.celrep.2014.03.020.

