## Supplementary figures and images for "Cut homeodomain transcription factor is a novel regulator of cortex glia morphogenesis and maintenance of neural niche"

### Supplementary Fig 1

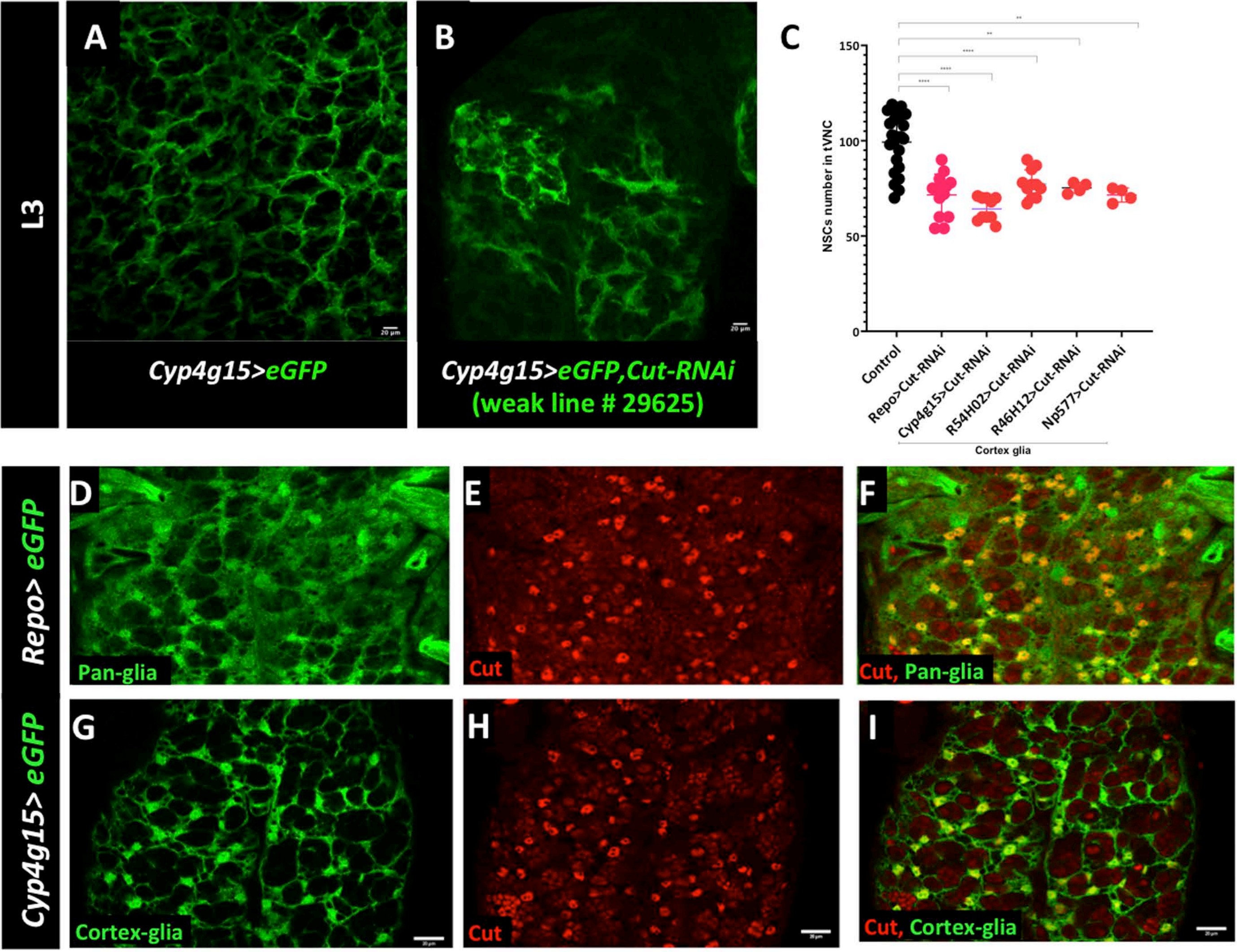


Supplementary Fig. 1

### Supplementary Fig 2

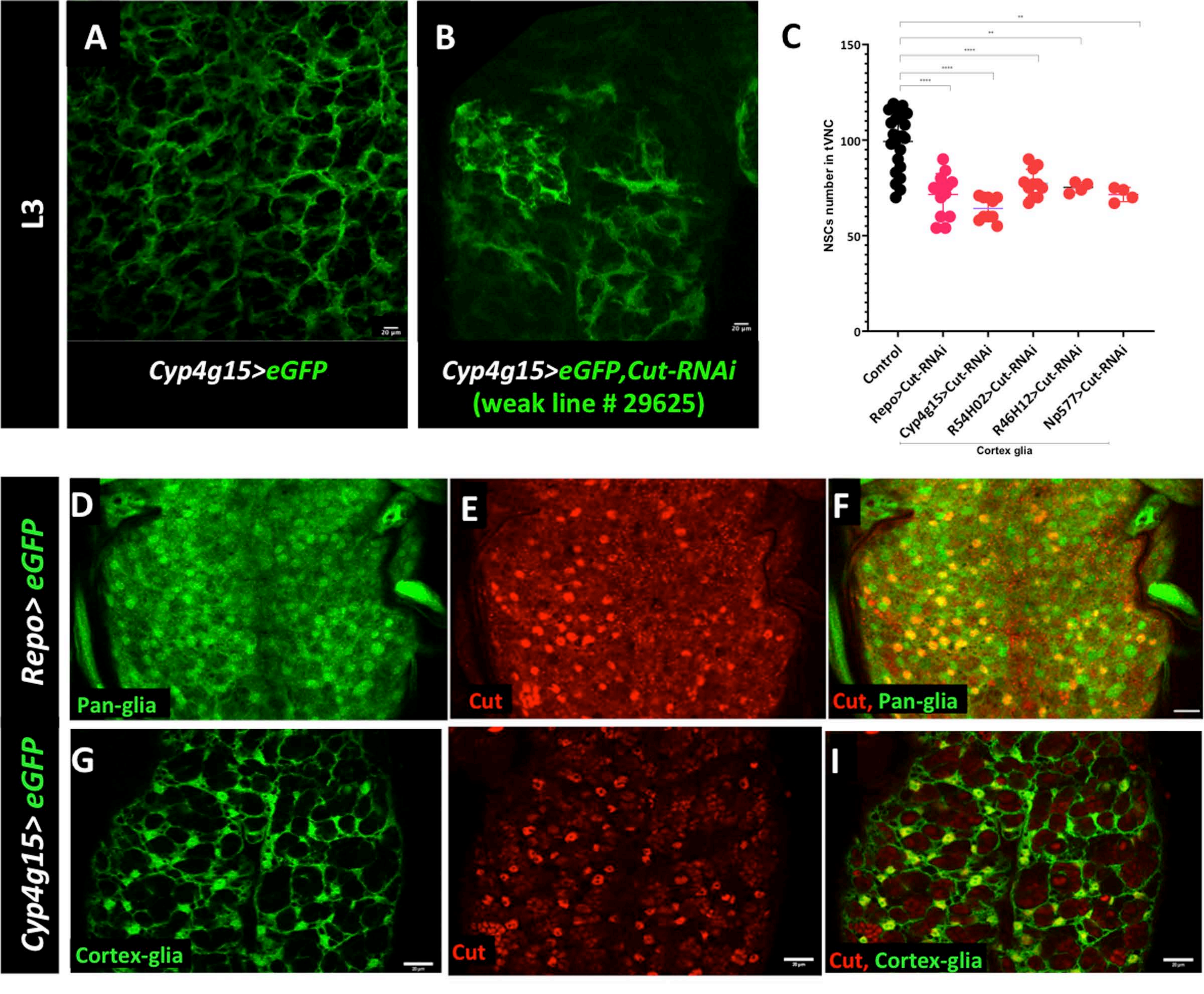


Supplementary Fig. 1

### Supplementary Fig 3

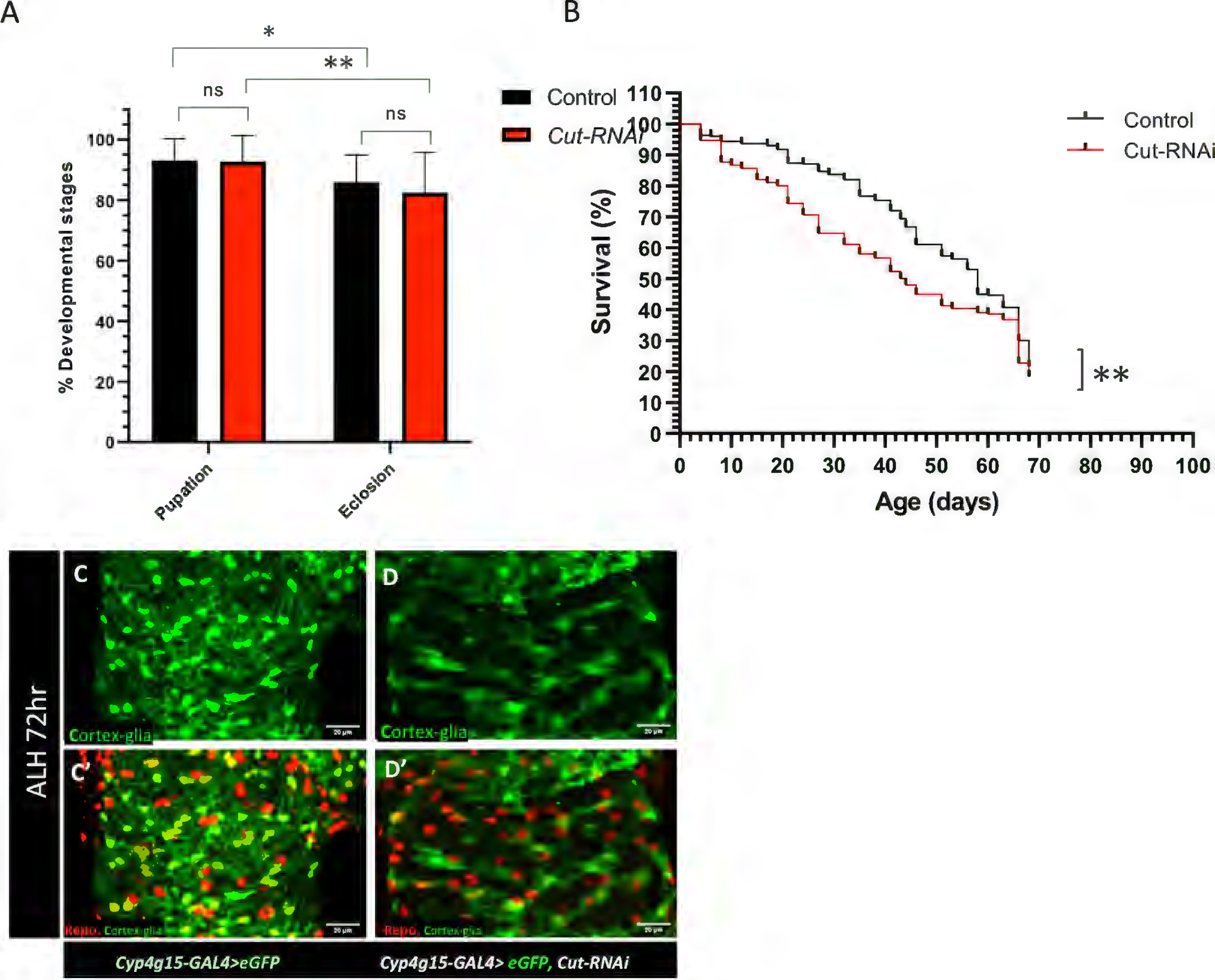


Supplementary Fig. 3
